# MEBOCOST: Metabolite-mediated Cell Communication Modeling by Single Cell Transcriptome

**DOI:** 10.1101/2022.05.30.494067

**Authors:** Rongbin Zheng, Yang Zhang, Tadataka Tsuji, Xinlei Gao, Allon Wagner, Nir Yosef, Hong Chen, Lili Zhang, Yu-Hua Tseng, Kaifu Chen

## Abstract

We developed MEBOCOST, a computational algorithm for quantitatively inferring metabolite-mediated intercellular communications using single cell RNA-seq data. The algorithm identifies cell-cell communications in which metabolites, such as lipids, are secreted by sender cells and traveled to interact with sensor proteins of receiver cells. The sensor proteins on receiver cell might be cell surface receptors, transporters across the cell membrane, or nuclear receptors. MEBOCOST relies on a comprehensive database of metabolite-sensor partners, which we manually curated from the literatures and other public sources. MEBOCOST defines sender and receiver cells for an extracellular metabolite based on the expression levels of the enzymes and sensors, respectively, thus identifies metabolite-sensor communications between the cells. Applying MEBOCOST to mouse brown adipose tissue (BAT) successfully recaptured known metabolite-mediated cell communications and further identified new communications. Additionally, MEBOCOST identified a set of BAT intercellular metabolite-sensor communications that was regulated by cold exposure of the mice. MEBOCOST will be useful to numerous researchers to investigate metabolite-mediated cell-cell communications in many biological and disease models. The MEBOCOST software is freely available at https://github.com/zhengrongbin/MEBOCOST.

## Introduction

Communication between cells, or cell-cell communication, is an integral part of cellular function in a human tissue. It is a critical process that maintains the functions and hemostasis of cells, organs, and intact systems^1^. Abnormal cell-cell communications are key contributors to many health anomalies such as obesity^2^, diabetes^3^, heart disease^4^, and cancer^5^. Communications between cells can be mediated by various types of molecules, e.g., proteins and metabolites. Protein-mediated cell-cell communications, e.g., those mediated by protein ligand-receptor pairs, have been the subject of many recent investigations based on single-cell RNA sequencing (scRNA-seq) and many robust algorithms^6-8^. Cell-cell metabolic reaction between cells is also frequently analyzed by inferring a metabolite generated by an enzyme in one cell and consumed as a substrate of another enzyme in a different cell^9^. For instance, free fatty acid (FFA) generated by lipase in adipocyte were reported to feed breast cancer cells, in which the FFA becomes substrate of acyl-CoA synthetases and is converted into Fatty Acel-CoA^10^. Several algorithms were recently reported to detect the generation and consumption of metabolites based on scRNA-seq data, thus enabled single-cell analysis of cell-cell metabolic reaction, e.g., COMPASS^11^, scFEA^12^, scFBA^13^. However, little computational resource is available to investigate metabolite-sensor communications, which is different from cell-cell metabolic reaction and cell-cell communications mediated by protein ligand-receptor pairs regarding the underlying molecular mechanism.

In a cell-cell metabolite-sensor communication, a metabolite generated by one cell travels to another cell, which has a sensor protein that binds the metabolite to trigger a signaling pathway^14, 15^. For instance, polyamine produced and secreted by EC was reported to be sensed by β-adrenergic receptor on the surface of white adipocyte to regulate adiposity^16^. Acetylcholine derived from B lymphocyte was recently reported to limit hematopoiesis by interacting with cholinergic α7 nicotinic receptor (Chrna7) on endothelial and mesenchymal stromal cells in bone marrow niche^17^. Li et al. reported that interactions between tumor histamine and histamine receptor H1 (HRH1) of macrophages rendered T cell dysfunctional^18^. In contrast to enzymes that consume the metabolite in the receiver cells of cell-cell metabolic reactions, sensor proteins in the receiver cells of cell-cell metabolite-sensor communications often do not consume the metabolite. Instead, sensor proteins often bind and release the metabolites to trigger and end cell signaling, respectively. Due to this mechanistic difference in the underlying biology, existing methods for analyzing cell-cell metabolic reaction are not applicable to analysis of metabolite-sensor communications. Meanwhile, most of current algorithms to analyze ligand-receptor communications were focused on protein or peptide ligands and thus designed in ways that do not support analysis of metabolite-sensor communications. Current major constraints on systematic analysis of cell-cell metabolite-sensor communications include the paucity of curated catalogue of reported metabolite-sensor partners and lack of robust method to detect active metabolite-sensor communications in a sample.

In this study, we addressed these most pressing constraints by developing a new bioinformatics technology, called MEBOCOST, which enables researchers to detect cell-cell metabolite-sensor communications by analyzing transcriptomes of single cells. We applied MEBOCOST to study cell-cell metabolite-sensor communications during thermogenesis, which is the heat-production process in a body and plays key roles in preventing many metabolic diseases such as the diabetes^19^. MEBOCOST successfully identified both known and new metabolite-sensor communications in brown adipose tissue (BAT) of mouse. It further identified cold-responsive communication events, which were reprogrammed in BAT under acute cold (2 days), chronic cold (7 days), room temperature, and thermal neutral conditions. By enabling systematic analysis of cell-cell metabolite-mediated communications, MEBOCOST will pave a new avenue for investigating metabolic signaling as the molecular basis of many development and disease processes.

## Results

### The MEBOCOST algorithm for detection of cell-cell metabolite-sensor communications

In metabolically active cells, thousands of expressed metabolic enzymes catalyze metabolic reactions to produce metabolites. Many metabolites can travel to extracellular space and function as signaling molecules. Some of these extracellular metabolites bind sensor proteins of spatially nearby cells. We termed the cells that secret the metabolites as sender cells and the cells that express the sensor proteins as receiver cells (Figure 1A). Therefore, interactions between these metabolites and sensor proteins mediate communications between the sender and receiver cells in a paracrine manner. They may also mediate autocrine when the sender cell and receiver cell are the same cell.

**Figure 1.**
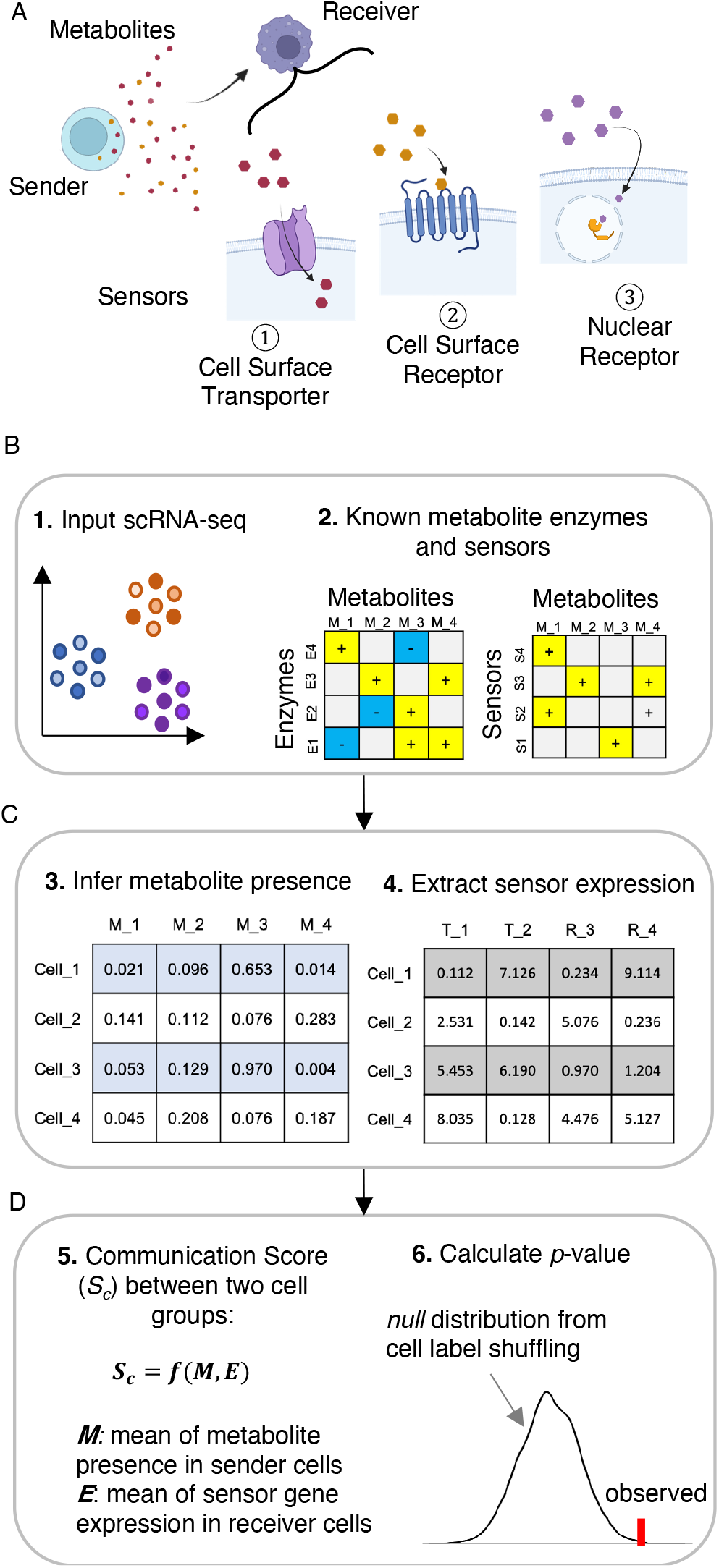
Cartoons to show flowchart in the MEBOCOST algorithm to preform metabolite-sensor communications between cells using scRNA-seq data. **A**. A cartoon to show that MEBOCOST predicts metabolite-mediated cell to cell communications, in which sender cells secret metabolites, and receiver cells receive metabolites through three mechanisms including cell surface transporter, cell surface receptor, and nuclear receptor. **B-D**. Cartoons to show the 6 steps to predict cell-cell metabolite-sensor communications in MEBOCOST. (1) The RNA expression data and cell type annotation obtained from scRNA-seq were taken as the input data to MEBOCOST. (2) A knowledgebase of enzymes and sensor proteins for metabolites is incorporated into the MEBOCOST algorithm. (3) The RNA expression values of metabolite enzymes were extracted from the scRNA-seq data to infer the presence of each metabolite based on the mean RNA expression of its enzyme genes. (4) The RNA expression values of the sensor proteins of metabolites were extracted from scRNA-seq data. (5) Calculation of communication score by taking the product of mean RNA expression of metabolic enzymes in the sender cell population and mean RNA expression of sensor proteins in the receiver cell population. (6) Shuffling single cell labels to generate a statistical *null* distribution to calculate *p* value for the communication score.

MEBOCOST integrates single cell RNA expression data and prior knowledge of extracellular metabolites, metabolic enzymes, and metabolite-sensor partners to detect metabolite-sensor communications (Figure 1B). To this end, MEBOCOST extracts RNA expression values of the enzymes from scRNA-seq data and infers the presence of metabolites in a cell based on the enzyme RNA expression (Figure 1C left, Supplementary Figure 1A-B). The enzyme genes of extracellular metabolites were collected from the Human Metabolome Database (HMDB)^20-23^, which provides annotation for 220,945 metabolites. Several algorithms have been reported to infer the presence of metabolite based on scRNA-Seq data. Therefore, we designed MEBOCOST to take the output of these algorithms as its input data. These algorithms include the scFBA^1413^, scFEA^1312^, and COMPASS^1211^ that each performed flux balance analysis in a unique way (Supplementary Figure 1C). Whereas flux balance analysis is popular for conventional metabolic analysis, the drawback is that it relies on some assumptions that might not be valid in many applications. Different assumptions will lead to different results. Also, flux balance analysis is computation-intensive and takes days or even weeks to do a genome-wide analysis of many single cells. In contrast, the mean RNA expression of ligand protein and receptor protein was frequently used to infer ligand-receptor communications, and is simple to calculate, biologically easy to interpret, and computationally efficient. Similarly, a gaussian mixture model was also reported to infer metabolite distribution based on expression of related enzymes^24^. It is yet unclear which methods might perform better when applied to infer presence of metabolite based on RNA-seq data. Therefore, we performed a comparison between several methods, including scFEA, COMPASS, and the geometric or arithmetic mean of RNA expression levels of related enzymes. The scFBA was excluded in this comparison because we developed MEBOCOST based on a free open access policy, whereas scFBA is implemented based on the commercial platform MATLAB. The gaussian mixture model is not included because no software or code was released yet to use this model. We used the four algorithms to analyze the CCLE dataset^25, 26^, which includes metabolomics data of 225 metabolites and genome-scale transcriptomics data from each of 928 cell lines. To measure the performance of each algorithm, we calculated Jaccard index for the overlap between metabolites inferred based on RNA expression and metabolites detected by metabolomics. Unexpectedly, despite being much simpler, we found that the arithmetic and geometric mean algorithms performed slightly better (greater Jaccard index value) than the COMPASS algorithms and as good as the scFEA algorithm (Supplementary Figure 1D). Further, the arithmetic and geometric mean algorithms are about 10 folds faster than scFEA and 1000 folds faster than COMPASS (Supplementary Figure 1E). Arithmetic mean performs slightly better than geometric mean. Therefore, we designed MEBOCOST to be compatible with all these four algorithms but set the arithmetic mean algorithm as the default option in MEBOCOST to infer metabolite presence.

Meanwhile, expression values of sensor genes were extracted from the scRNA-seq data (Figure 1C right). Next, the metabolite presence score calculated by MEBOCOST, scFEA, or COMPASS as the estimated likelihood of metabolite presence, along with sensor gene expression value, were averaged per cell cluster defined in advance by users based on scRNA-seq analysis. For each metabolite-sensor communication event, a communication score will be calculated by taking the product of the average metabolite presence score in the sender cell group and the average expression of the sensor protein in the receiver cell group (Figure 1D left). Such calculations were performed for each metabolite-sensor partner and each pair of cell clusters. A higher communication score represents a higher likelihood of the metabolite-sensor communication between the associated two cell clusters. Finally, the statistical significance of a communication score was evaluated based on permutation test. For each metabolite-sensor communication, MEBOCOST generated a null statistical distribution for the communication score by shuffling the cell labels of all cells in the scRNA-seq data (shuffle 1000 times by default). The *p* value for the communication score was determined based on the null distribution. The *p* values of individual communications were further corrected to calculate false discovery rate (FDR) using the Benjamini-Hochberg’s method^27^.

### A curated knowledge repository of enzyme-metabolite-sensor partners

To collect the extracellular metabolites and their enzymes, we started from the 157 and 2,597 unique metabolites covered by scFEA and COMPASS, respectively (Figure 2A steps a-b). To focus on a subset of well-annotated metabolites, we retrieved the metabolites that could be mapped to those in the HMDB and further removed redundant metabolites that represent different names of the same metabolite. This resulted in 1,240 metabolites with unique accession number from the HMDB (Figure 2A step c). As our interest is in cell-cell communications, we further narrowed down to 910 metabolites annotated as could be in extracellular space, blood, or cerebrospinal fluid (Figure 2A step d). We defined these metabolites as extracellular metabolites in this study. To infer metabolite presence based on enzyme RNA expression, we next selected a subset of 441 metabolites that have annotated upstream enzymes in metabolic reactions (Figure 2A step e).

**Figure 2.**
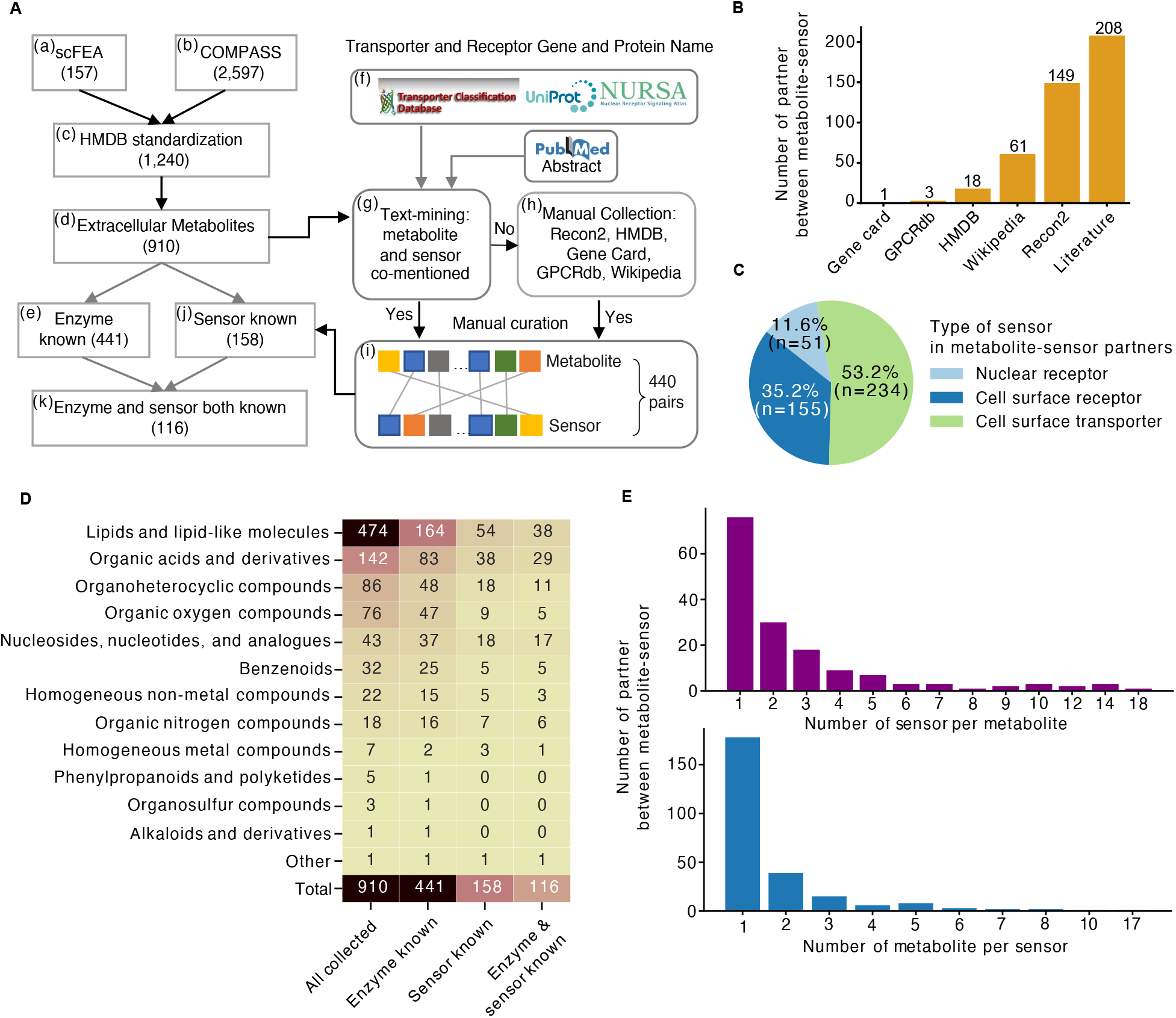
The collection of extracellular metabolites, enzymes, and sensor proteins. **A**. The pipeline to collect extracellular metabolites and their enzymes as well as sensors. **B**. Bar plot to show number of metabolite-sensor partners collected from different sources. **C**. A pie chart to show the number of each type of sensor proteins in the collected metabolite-sensor partners. **D**. A heatmap to show the number of each type of metabolites in the collected metabolite-sensor partners. **E**. Number of metabolite-sensor partners plotted against number of sensors per metabolite (top panel) and number of metabolites per sensor (bottom panel).

Although metabolite-sensor partners have been reported by many individual studies, few efforts were taken to thoroughly collect and curate the catalogue of reported metabolite-sensor partners. To this end, we first developed a computational pipeline for comprehensive literature mining. We first retrieved the 233 transporters from the Transporter Classification Database^28-32^, the 1,522 cell surface receptor proteins from the UniProt database^33^, and the 48 nuclear receptors from the Nuclear Receptor Signaling Atlas (NURSA) database^34^ (Figure 2A step f). Next, a text-mining approach was developed to parse the metabolite-sensor partners from the abstracts of publications in the PubMed database. Briefly, an abstract will be selected if any pair of metabolite and sensor were co-mentioned in a sentence from the abstract (Figure 2A step g). The obtained metabolite-sensor partners together with the corresponding abstracts were then subjected to three rounds of manual curation. At least three curators read each abstract independently and filter out the metabolite-sensor partners for which the evidence in the abstracts was insufficient. We also searched for annotated metabolite-sensor partners in five annotation databases: Recon2^35^, HMDB^20-23^, GeneCards^36^, GPCRdb^37, 38^, and the Nuclear Receptor page of the Wikipedia site (Figure 2A step h). Altogether, combing the results from these procedures result in 440 metabolite-sensor partners in total (Figure 2A step i), of which 208, 149, 61, and 22 partners were collected from literature, Recon2, Wikipedia, and other sources, respectively (Figure 2B). Among these partners, 53.2% was metabolite-transporter partners, 35.2% was metabolite-cell surface receptor partners, and 11.6% was metabolite-nuclear receptor partners (Figure 2C). We retrieved the 158 metabolites in these partners (Figure 2A step j), and further narrowed down to the 116 that also have known enzyme (Figure 2A step k). Of these 116 metabolites, most (33%) are lipids and lipid-like molecules (Figure 2D). Organic acids and derivatives also constitute a major proportion (25%). Other major categories include Nucleosides, nucleotides, and analogues (15%), Organoheterocyclic compounds (9%), along with 10 other minor types. Among the collected metabolite-sensor partners, most metabolites each have one sensor, while some metabolites each have multiple sensors (Figure 2E). Also, most sensors each have one metabolite, while some sensors each have multiple metabolites. This database of metabolite-sensor partners represents a rich research resource for researchers to investigate metabolite-sensor communications.

### Great scalability and robust performance of MEBOCOST

To evaluate the usage of computing resource, we tested MEBOCOST based on a scRNA-seq dataset generated for stromal vascular fraction of brown adipose tissue (BAT) from mouse housed at cold temperature (4 °C) for 2 days (Cold2)^39^. The data includes 33,470 cells in total. We applied MEBOCOST on the origional data as well as a series of down-sampled data. We found that the time and computer memory taken by MEBOCOST had a linear correlation with the data size. The time to run MEBOCOST ranged between 9 and 7 minutes when the number of total cells was sampled down from 33,470 to 11,157 (Figure 3A), where the peak memory usage ranged from 2.5 to 1.5Gb (Figure 3B). We also evaluated the effect of sequencing depth on the performance of MEBOCOST. The result indicated that the number of detected communications showed little change even after the total cell numbers were sampled down to keep only 30% of the original data (Figure 3C). To compare the similarity of detected communications between down-sampled datasets and original dataset, we introduced two measurements including “recaptured over total detected” and “recaptured over originally detected”. In the recaptured over total detected, the number of communications detected in both the down-sampled and original datasets were divided by the number of communications detected in the down-sampled dataset. In the recaptured over originally detected, the number of communications detected in both the down-sampled and original datasets were divided by the number of communications detected in the original dataset. We observed that both “recaptured over total detected” and “recaptured over original” were still greater than 80% even when the number of total cells was sampled down to 30% of the original data (Figure 3D).

**Figure 3.**
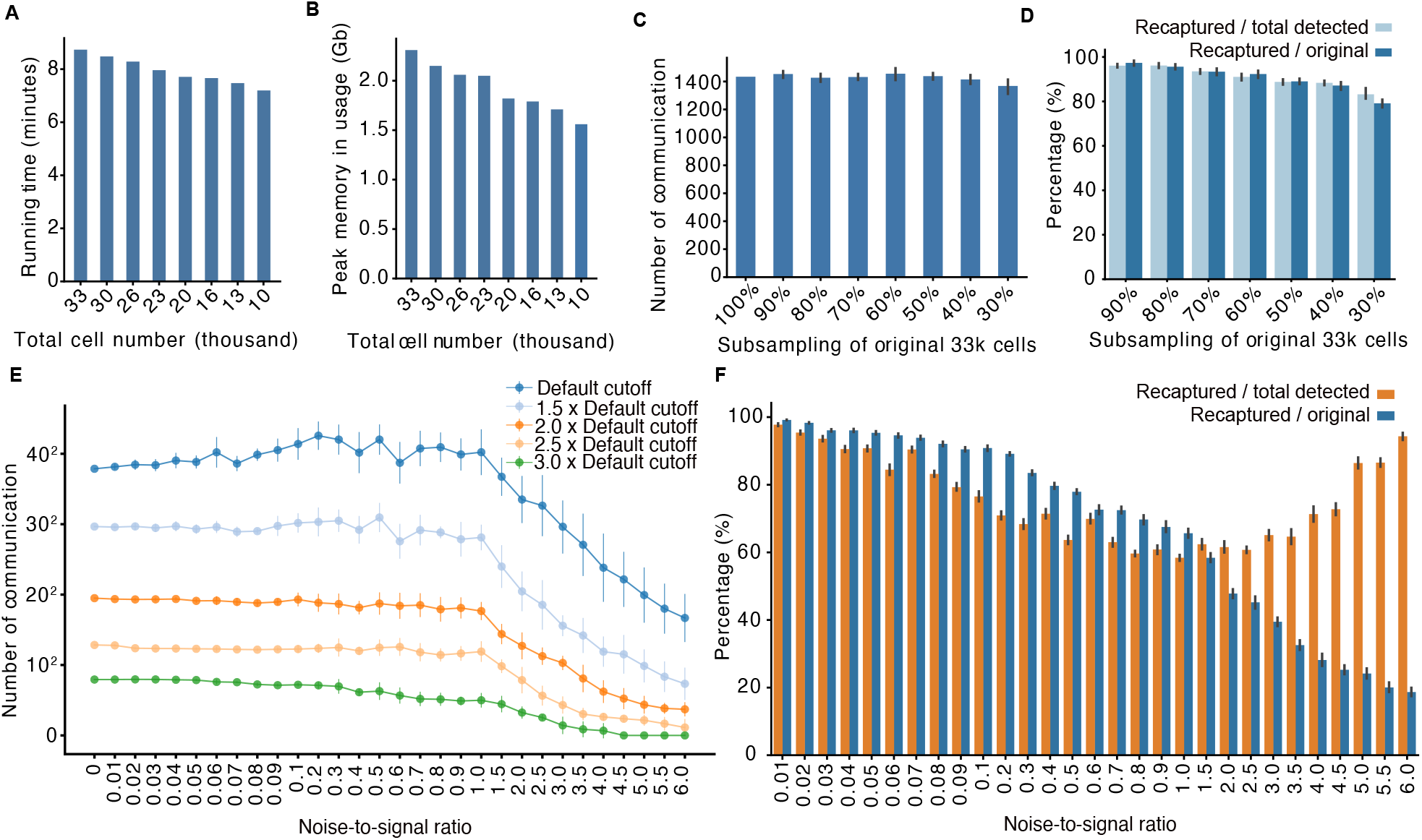
Robust performance and great scalability of MEBOCOST. **A-D**. Bar plots to show the running time (A), peak memory usage (B), number of detected communications (C), and recapture rate (D) of MEBOCOST when applied to a series of down sampled scRNA-Seq datasets. The error bar showed range of the results from 10 repeats of down sampling experiments. **E-F**. The number of detected communications (E) and recapture rate (F) plotted against simulated noise-to-signal ratio after applying MEBOCOST to a series of simulated scRNA-seq data. The original sequenced reads were recognized as the signal, whereas noise was simulated by adding random reads to the scRNA-seq data. The error bar showed the range of results from 10 repeats of the simulation experiments. All down sampling and simulation experiments were performed based on mice BAT data from the Cold2 condition.

We next tested the tolerance of MEBOCOST to sequencing noise. To this end, we generated a set of simulated noising dataset with a series of noise rate by adding random reads to the Cold2 scRNA-seq data. Next, we applied MEBOCOST on these datasets and tested the ability to recapture the communications detected in the original Cold2 scRNA-seq data (Figure 3E). The results showed that the number of detected communications reduced substantially when noise read number increased from 1 to 6 folds of the original scRNA-seq read number (noise-to-signal ratio from 1 to 6) (Figure 3E). However, the recaptured over total detected communications increased substantially, whereas the recaptured over originally detected decreased (Figure 3F). These results suggest that MEBOCOST is unlikely to detect communications from noise signal. In datasets with noise-to-signal ratio less than 1, the number of detected communications remained stable (Figure 3E). Although the recaptured over total detected and recaptured over original detected both decreased slightly in response to the moderate increase of the noise-to-signal ratio from 0.01 to 1, both measurement remains higher than 60% (Figure 2F). Therefore, the performance of MEBOCOST is resistant to moderate noise-to-signal ratio lower than 1. Also, the fluctuation of number of detected communications at different noise-to-signal ratios appeared to be moderate and can be further decreased when the cutoff to define communications was set to be more stringent (Figure 2E). Altogether, the subsampling and noise simulation strategies both demonstrated that MEBOCOST is a robust algorithm for predicting metabolite-sensor communications from scRNA-seq data.

### MEBOCOST recaptured known and identified new metabolic communications in BAT

BAT is a metabolically active tissue and is specialized to dissipate chemical energy in the form of heat in thermogenesis. Cold exposure has been known as a strong activator for BAT^40-42^. It is well recognized that cold exposure promotes not only thermogenesis but also the process of adiposity or differentiation of brown adipocytes^43^. The process of brown adipocyte differentiation can be regulated by metabolites in BAT^44, 45^. We speculated that different cell types in BAT can communicate through metabolites and the intercellular communications with adipocytes regulate the differentiation and function of adipocytes. Therefore, for comparison with the Cold2 data, we further analyzed scRNA-seq data generated for BAT stromal vascular fraction of mouse housed with thermoneutral (TN) temperature (30 °C for a week), room temperature (RT, 22 °C), and chronic cold (Cold7) temperature (4 °C for 7days)^39^. The data from the four conditions include 107,679 high-quality cells and 468 million reads from twenty cell types (Supplementary Figure 2A-E). These include adipocytes, Schwann cells, vascular cells, immune cells, etc. Mature adipocytes are enriched with lipid droplets and tend to be excluded from the stromal vascular fraction, thus tend to be not captured by the scRNA-seq experiment. Therefore, most adipocytes in this scRNA-seq dataset would be differentiating adipocytes.

We first analyzed the BAT scRNA-seq data of the Cold2 condition, as it contains more adipocytes when compared to the other three conditions (Supplementary Figure 2F). In total, 1,434 metabolite-sensor communications were detected in the Cold2 BAT based on an FDR cutoff 0.05 (Figure 4A). In addition to paracrine communications between cell types, autocrine communications of each cell type were also observed for many cell types such as adipocytes, endothelial cells (EC), lymphatic EC, basophils/eosinophils, ILC2s, etc. Among all communications, adipocytes showed the largest number of communications compared to other cell types (Figure 4B). Many of the communications between adipocytes and other cell types showed higher overall score compared to those between other cell types (Figure 4A). Notably, all cell types each can be both senders and receivers in the metabolite-sensor communications in BAT. 2.4-fold greater number of communication events were observed for adipocytes as sender cells (177 events) than as receiver cells (72 events) (Figure 4B). In addition, of all the communications sent to adipocytes, autocrine of adipocytes happened more frequently than paracrine (Figure 4C). Examples of such metabolite-sensor partners include the Myristic acid ∼ Slc27a2, L-Glutamine ∼ Slc1a5, L-Glutamine ∼ Slc38a2, Vitamin A ∼ Rbp4, Eicosapentaenoic acid ∼ Ffar4, and Docosahexaenoic acid ∼ Ffar4 (Figure 4D-E). Some of the metabolites, sensors, and metabolite-sensor partners have been reported to be involved in regulating the function of BAT. For example, the eicosapentaenoic acid (EPA) and docosahexaenoic acid (DHA), a type of omega-3 fatty acid, were reported to play an important role in brown adipocyte differentiation and thermogenesis^46^. Particularly, Jiyoung Kim and colleagues reported that EPA can potentiate brown adipocyte thermogenesis dependent on Ffar4^47^. The critical role of vitamin A transport has also been demonstrated in adipose tissue browning and thermogenesis upon cold exposure^48, 49^. Such importance was proven by knocking out Rbp, which is a gene encoding the vitamin A transporter^49^. Glutamine was reported as a major source of *de novo* fatty acid synthesis in a brown adipocyte cell line^50^, and the utilization of glutamine was reported to be enhanced in brown adipose tissue by acute cold exposure^51^. Meanwhile, some metabolite-sensor communications were newly identified by MEBOCOST in this study. Among those, myristic acid and Slc27a2 were particularly noticed since it had the highest communication score among the detected adipocyte autocrine communications (Figure 4D). Interestingly, myristic acid was specifically and highly enriched in adipocytes when compared to other cell types in Cold2 BAT (Figure 4F top panel). Similarly, the mRNA expression of Scl27a2, a transporter for myristic acid, was also expressed specifically and higher in the adipocytes than in other cell types (Figure 4F bottom panel). These results showed that MEBOCOST successfully recaptured known and further discovered new metabolite-sensor communications for adipocyte autocrine.

**Figure 4.**
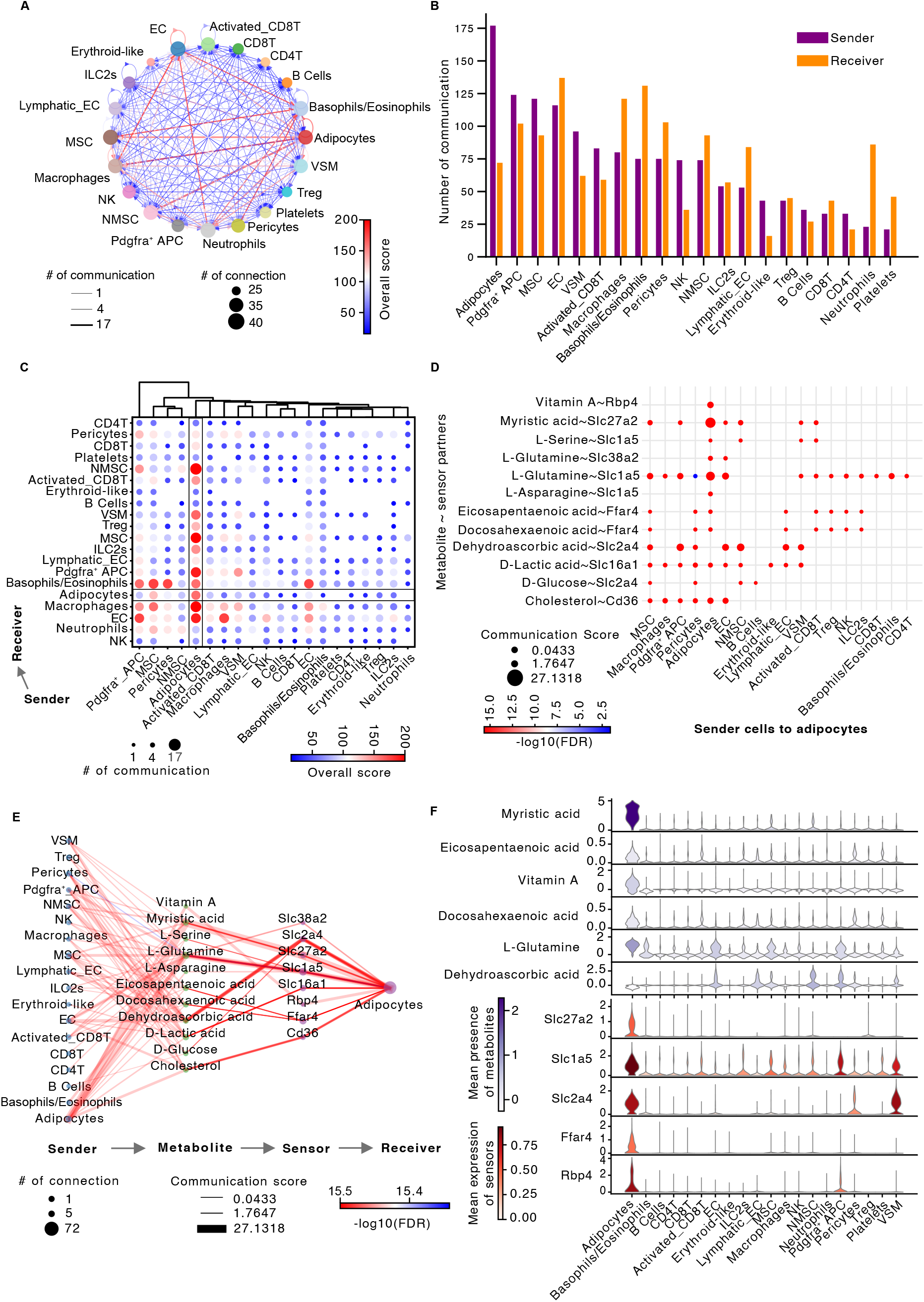
MEBOCOST recaptured known and revealed new metabolite-sensor communications between cells in brown adipose tissue mice exposed to cold for 2-days. **A**. a network plot showing cell-to-cell metabolite-sensor communications detected by MEBOCOST for the Cold2 BAT scRNA-Seq data. Each dot was a cell type. The size of dot for each cell type represented the number of communications with the other cell types. Each arrow line represented the communication from a sender cell type to a receiver cell type. The line width indicated the number of metabolite-sensor communications between the sender and receiver cell types. Line color showed the overall communication score which was calculated by the sum of - log10(FDR) of all metabolite-sensor communications between the sender and receiver types. **B**. Bar plot to show the number of detected communications with each cell type as sender cells or receiver cells. **C**, A dot plot showing the communications between each pair of cell types. The dot size indicated the number of metabolite-sensor communications between a pair of sender and receiver cell types. The color of the dot represented the overall score of communications between the sender and receiver cell types, with the score calculated in the same way as in the panel A. The black lines highlighted adipocyte associated communications. **D**. A dot plot showing metabolite-sensor communications send to adipocytes from adipocytes themselves and other cell types. The rows represent metabolite-sensor partners, the columns represent the sender cell types in the communications with adipocytes. The dot size indicated the communication score calculated by MEBOCOST. The color of the dot was the −log10(FDR) for the metabolite-sensor communication score. **E**. A flow diagram showing the information flow of metabolite-sensor communications from sender cell types to receiver cell types. The size of dots represented the number of connections in the diagram. The lines connect the sender cell, metabolite, sensor, and receiver cell. Line width represented the metabolite-sensor communication score. Line color indicated the −log10(FDR) of the communication score. **F**. Violin plots showing the RNA expression of enzymes (top panel) and sensor proteins (bottom panel) of metabolites in individual cell types of brown adipose tissue.

In addition to autocrine, adipocytes received many paracrine-style metabolite-sensor communications from other cell types in the brown adipose tissue. The most frequent sender cell types in the communications with adipocytes include myelinating Schwann cell (MSC), pericytes, activated CD8^+^ T cells (CD8T), vascular smooth muscle cells (VSM), non-myelinating Schwann cell (NMSC), Pdgfra^+^ adipocyte progenitor cells (APC), and lymphatic EC, etc (Figure 4C). Among these sender cells, the communications between vascular system and adipocytes were well known^52^. For instance, Jennifer H Hammel and Evangelia Bellas reported that EC-adipocyte crosstalk can improve adipocyte browning, as proven based on an EC-adipocyte co-culture system^53^. Meanwhile, some senders were newly detected by MEBOCOST. For instance, CD8T cells, a subtype of CD8T cells expressing cytotoxic markers, were much less reported regarding metabolite-sensor communication with adipocytes in BAT. Interestingly, 5 communications from activated CD8T cells to adipocytes were detected in Cold2 BAT, while only one communication were detected for naïve CD8 T cells (Figure 4D). Among all metabolite-sensor partners, the communication score of dehydroascorbic acid and Slc2a4 were among the highest (Figure 4D-E). Such communications mediated by dehydroascorbic acid and Slc2a4 were detected between MSC and Adipocytes, Pdgfra^+^ APC and Adipocytes, Pericytes and Adipocytes, NMSC and Adipocytes, EC, and Adipocytes, VSM and Adipocytes, as well as Lymphatic EC and Adipocytes. Moreover, the metabolite presence inference showed that dehydroascorbic acid was highly enriched in those sender cells including Pdgfra^+^ APC, NMSC, EC, and VSM when compared to other cell types (Figure 4F top panel). Meanwhile, we found that Slc2a4 was also highly and specifically expressed in brown adipocytes compared to most of other cell types (Figure 4F bottom panel), indicating that those communications discussed above might function importantly for adipocytes. In summary, the analysis of MEBOCOST based on Cold2 BAT scRNA-seq data showed that the brown adipose resident cells frequently communicate with other cells in the microenvironment through metabolite-sensor partners.

### MEBOCOST identified cold-sensitive metabolite-sensor communications in BAT

We next performed comparative analysis of metabolite-sensor communications in BAT among the four conditions including TN, RT, Cold2, and Cold7. We first used MEBOCOST to analyze BAT scRNA-seq data from each of the four conditions separately. The number of detected metabolite-sensor communications increased during thermogenesis in response to cold exposure, especially by the chronic cold exposure for 7 days (Figure 5A). In total, 1,330, 1,314, 1,434, and 1,649 communication events were detected in BAT under the TN, RT, Cold2, and Cold7 conditions, respectively. Many metabolites and sensors appeared to be cell type- and condition-specific (Figure 5B, C). Furthermore, the composition of communications between cell types in BAT was also dramatically changed by cold exposure in mice. For instance, the Pericytes received the largest number of communications from other cell types under the TN and RT conditions (Supplementary Figure 3A-D), while EC and basophils/eosinophils were the cell types receiving the largest number of communications under the Cold2 and Cold7 conditions, respectively (Figure 4A-B, Supplementary Figure 3E-F). Similar changes were observed for sender cells. The MSC was the most frequently used sender cell type in TN, while macrophages, adipocytes, and EC were identified to send the greatest number of metabolites to other cell types in RT, Cold2, and Cold7, respectively (Figure 4A-B, Supplementary Figure 3). These results suggested that the intercellular metabolite-sensor communications in mouse BAT were regulated by environment temperatures.

**Figure 5.**
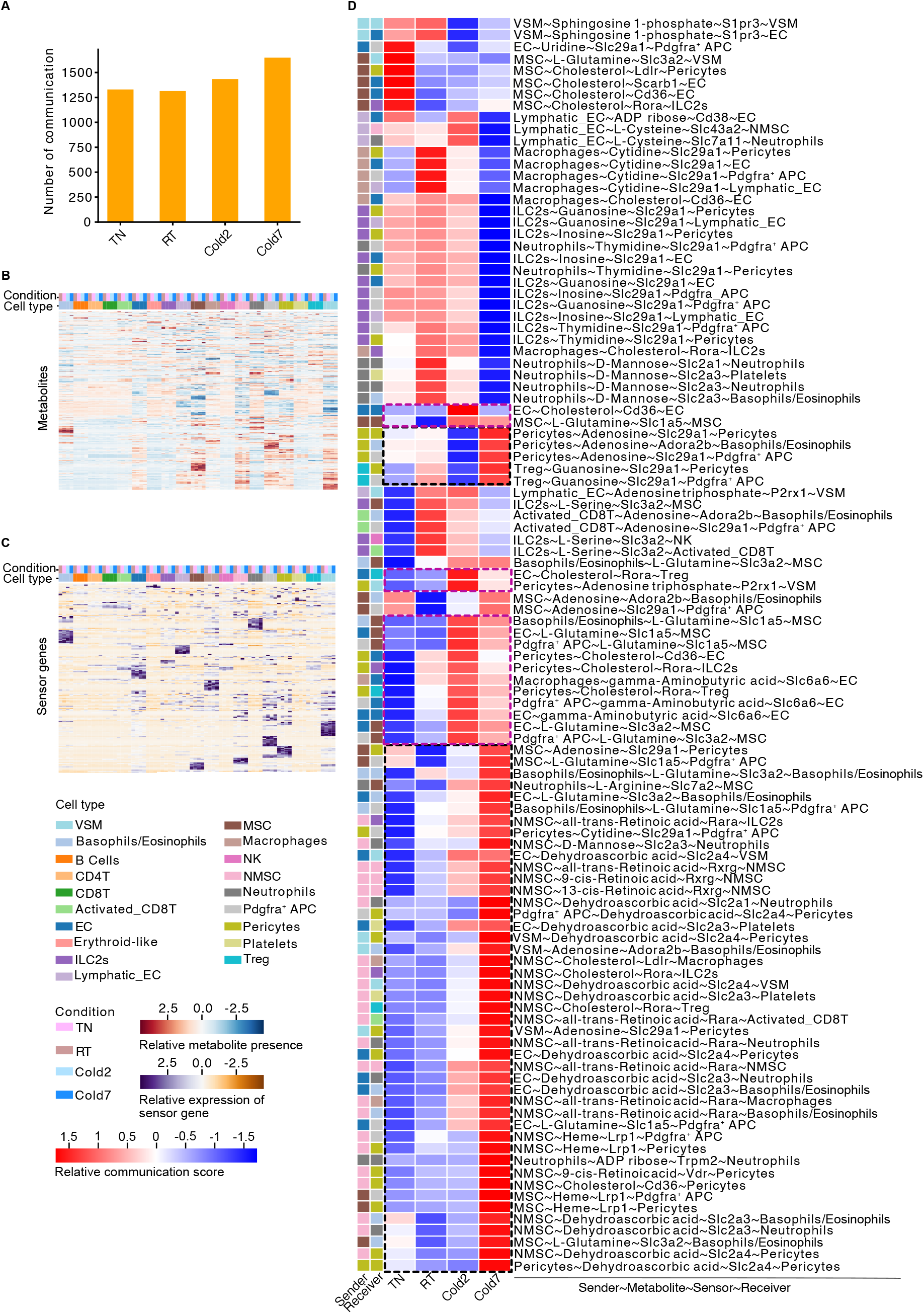
MEBOCOST revealed temperature-sensitive metabolite-sensor communications in brown adipose tissue. **A**. A bar plot showing the total number of communications detected by MEBOCOST in BAT scRNA-seq data from mice under the TN (30 °C for a week), RT (room temperature), Cold2 (4 °C for 2 days), and Cold7 (4 °C for 7 days) conditions. **B-C**. A heatmap to show the estimated metabolite presence (B) and RNA expression of metabolite sensors (C). Columns were cell types in the four conditions and rows were metabolites. **D**, A heatmap showing detected metabolite-sensor communications that were sensitive to cold exposure. Each row represented one cell-cell metabolite-sensor communication. A communication was defined as a combination of sender cell type, a metabolite, a sensor protein, and a receiver cell type. The purple and black dashed boxes highlighted communications that were increased by cold exposure.

Next, to identify the temperature-sensitive metabolite-sensor communications, all cells from the four conditions were pooled for MEBOCOST analysis and then grouped by cell types and conditions. With the detected communications from the four conditions, index of dispersion (IOD) of the communication scores was calculated to characterize the sensitivity of communications to cold exposure. The temperature-sensitive communications were identified as with the 5% greatest variation (IOD > 0.33) of communication scores across conditions (Figure 5D). Among these, we observed 65 communications with increased communication score under any of the two types of cold exposure including both Cold2 and Cold7 (Figure 5D, black and purple dashed box labeled). 50 communications were strongly and specifically increased by chronic cold exposure for 7 days in Cold7 (Figure 5D, black dashed box labeled). Among the specifically increased communications in Cold7, NMSC was the major sender cell type compared to other cell types and contributed to 46% (23 out of 50) of the communications. This result indicates that chronic cold exposure for 7 days may increase the secretion of signaling metabolites from NMSC and increase its effects on other adipose-resident cells. Altogether, MEBOCOST enabled detection of condition-specific cell-cell metabolite-sensor communications in BAT.

## Discussion

Cell-cell communication is a fundamental mechanism that coordinates cellular activities in development and disease^6^. In addition to protein ligands, metabolites are another major type of signaling molecules to mediate cell-cell communications, which are studied extensively based on experimental approaches. However, few computational tools were appliable to study metabolite-based intercellular communications based on single cell RNA-seq data. Motivated by the fundamental importance of metabolic communications in development and diseases, we developed MEBOCOST to fill up the technology gap and infer intercellular metabolite-sensor communications using scRNA-seq. To this end, we first constructed a metabolite-sensor partner database, which covered sensor types including cell surface receptor, cell surface transporter, and nuclear receptor. We developed a computational pipeline followed by manual curation for a thorough literature mining to collect reported metabolite-sensor partners from literature. The collected metabolite-sensor partners provided a new foundation of knowledge for investigating cell-cell metabolite-sensor communications. Having those partners, communication scores were calculated based on the expression of metabolic enzymes and sensors for each pair of cell types and each metabolite-sensor partner to characterize the communication likelihood. By applying to a BAT scRNA-seq datasets, MEBOCOST successfully recaptured known metabolite-sensor communications and further uncovered new communications. As a versatile and easy-to-use Python-based package, MEBOCOST will enable researchers to analyze cell-cell metabolite-sensor communications based on scRNA-seq data in numerous human and mouse samples.

Two recently released algorithms, including the CellPhoneDBv4^54^ and scConnect^55^, could perform analysis of cell-cell communications mediated by some small molecules. Likely because these two algorithms were designed mainly for analysis of protein ligand-receptor analysis, only 199 small molecule-receptor partners and 8 neurotransmitter-receptor partners were included respectively. In contrast, MEBOCOST included transporters in addition to receptors of metabolites and currently covered 440 partners, thus is much more comprehensive. A recent set of human genome-scale metabolic models (GEMs) contain at least 4,164 unique metabolites^*56*^. Our preliminary work started with 1240 unique metabolites that scFEA and COMPASS have covered. The default algorithm in MEBOCOST to infer metabolite presence does not have to rely on scFEA and COMPASS. Therefore, as we continue to further curate known metabolite-sensor pairs for the additional metabolites using the pipeline that we have established, we predict that MEBOCOST will ultimately cover about 3 folds more metabolites. CellPhoneDBv4 utilized the gene expression of the last enzyme in the biosynthesis pathway to represent a small molecue in the calculation of communication score. scConnect used several key enzymes of the biosynthesis and uptake pathways in their calculations. However, many metabolites or small molecules are participated in complex metabolism which includes both production and consumption for the metabolites. CellPhoneDBv4 and scConnect both did not take consumptive reactions of a metabolite into consideration when inferring metabolite presence by scRNA-seq data. In contrast to these two algorithms, MEBOCOST algorithm considered both the generation and consumption reaction of a metabolite. Further, MEBOCOST was designed to be compatible with mature algorithms that infer single-cell metabolite presence based on flux balance analysis, including the scFEA and COMPASS algorithms.

To the best of our knowledge, MEBOCOST is the first computational algorithm that is specifically dedicated for cell-cell metabolite-sensor communications. By continuing to develop new algorithms and functions, the capability of MEBOCOST will be further enhanced through multiple major aspects in the future. The number of extracellular metabolites can be substantially increased by integrating more metabolite resources such as HMDB, Recon2, Metabolic Atlas, etc. The text-mining pipeline to improve the recognition of metabolite-sensor partners will also be further optimized. For example, we will enable the pipeline to use synonymous of metabolites and sensors to parse the metabolite-sensor partners from PubMed abstract. Also, due to the complicated biological mechanism of cellular communication, taking more information into consideration in communication prediction will increase the performance of the algorithm. Cell surface transporter or receptor for metabolites might function by a protein complex. For example, SLC7A11 is a cell surface transporter for Cysteine curated in our database, and SLC3A2 is an extra need for forming the SLC7A11 mediated metabolite-sensor communications^57^. Therefore, it might be useful to improve MEBOCOST by incorporating the protein complex information of sensors into the calculation of communication score. Considering that the relation between metabolite abundance and RNA expression level of metabolic enzymes are often nonlinear, we do not expect quantitatively calculate the abundance of metabolite by our current algorithm. We instead have aimed at inferring the presence of metabolite based on scRNA-Seq data and interpret it as a qualitative result. A more sophisticated algorithm to quantitatively predict metabolite abundance, such as machine learning model, might be helpful to further improve accuracy of metabolite-mediated cell communication analysis. Finally, MEBOCOST infers potential metabolite-sensor communications using single cell transcriptomics data without considering the spatial proximity of the cells. Recently, spatial transcriptomics has been successfully utilized to improve the prediction of protein-based cell-cell interaction by many other computational tools^5, 7, 8, 54^, demonstrating the informative role of spatial information in the cell-cell communication prediction. Although a robust performance has been observed in the current MEBOCOST, we believe that it will have the potential to provide a more comprehensive view of metabolite-sensor communications by combining spatial distribution of the cells into consideration.

## Methods and Materials

### Software and package version

Cellranger 6.1.2, scFEA 1.1.2, COMPASS (https://github.com/YosefLab/Compass, downloaded on October 19^th^, 2021), Python 3.8, and python packages including pandas 1.4.1, scipy 1.8.0, scanpy 1.8.2, matplotlib 3.5.1, seaborn 0.11.2, adjustText 0.7.3, network 2.7, jupyter 1.0.0, MEBOCOST v1.0.2.

### Collection of extracellular metabolites and related enzyme genes

MEBOCOST analyzes metabolites that are predictable from gene expression data. scFEA and COMPASS are two software for metabolic flux-balance analysis in scRNA-seq data. The metabolites covered by this two software were analyzed. Specifically, the metabolite names and their annotation information in scFEA were downloaded from https://github.com/changwn/scFEA/blob/master/data/Human_M168_information.symbols.csv. The metabolite name and annotation in COMPASS were downloaded from https://github.com/YosefLab/Compass/blob/master/compass/Resources/Recon2_export/met_md.csv. We noticed that HMDB at https://hmdb.ca/ is a comprehensive database that include very detailed annotation for 220,945 metabolites. Therefore, we decided to standardize the collected metabolite names from both scFEA and COMPASS into the annotation provided by HMDB. To do this, we focused on the metabolites for which the HMDB accession number was known. HMDB accession numbers of metabolites can be directly accessed in COMPASS, while only KEGG accessions were provided for scFEA metabolites. To map those scFEA metabolites into HMDB annotation, a parser script was developed to covert KEGG compound accession number into HMDB accession number. Taking Acetyl-CoA as an example, C00024 is the KEGG compound accession number, and the KEGG page of C00024 is at https://www.genome.jp/entry/C00024. The related annotation in HMDB for Acetyl-CoA can be further found by hyperlinks at the “All links” session. In this specific case, the HMDB accession number for Acetyl-CoA can be collected from https://www.genome.jp/dbget-bin/get_linkdb?-t+hmdb+cpd:C00024. We applied the strategy for all the metabolites from scFEA to find their the HMDB accession numbers. For those metabolites that were successfully mapped to HMDB, biological location annotation, including cellular locations and biospecimen locations, were extracted. To focus on potential intercellular signaling metabolites, the metabolites in “extracellular space”, “blood”, or “cerebrospinal fluid” were retained and named as “extracellular metabolites”. Meanwhile, other basic annotation information for those metabolites were also collected, such as synonyms, metabolite class, and protein associations.

To collect metabolite enzymes, we focused on the extracellular metabolites collected by the procedures mentioned above. A parser script was developed specifically to collect the reaction and related enzymes of extracellular metabolites in HMDB. Briefly, the webpage of metabolite in HMDB was visited based on the given HMDB accession number, and the annotation in the “Enzymes” sessions was collected. Taking D-Lactic acid as an example, the annotation page of D-Lactic acid can be retrieved by https://hmdb.ca/metabolites/HMDB0001311 where HMDB0000171 is the HDMB accession number. In the “Enzymes” tab, the reactions such as “S-Lactoylglutathione + Water → Glutathione + D-Lactic acid” and the related gene name (e.g. HAGH) for the reactions were collected. Only complete reactions containing subtract metabolite names and product metabolite names were included in the collection. Such procedures produce a list of reactions as well as the corresponding enzyme gene names and metabolite names. The collection of reaction and related enzymes were used for the inference of metabolite presence.

### Collection of metabolite-sensor partners

The detection of metabolite-based intercellular communications relied on the prior knowledge of metabolite-sensor partners. The sensor proteins in this study mainly include three types, namely, cell surface transporter, cell surface receptor, and nuclear receptors. To collect known pairs of metabolite-sensor partners, a workflow that combines computational text-mining and manual collection was developed.

In the text-mining part, a list of metabolites and a list of sensor gene names were needed. For metabolites, we focused on the extracellular metabolites. As one category of sensor proteins, we obtained cell surface receptors from two major sources. Firstly, we downloaded a list of receptors from NicheNet^58^, a protein ligand-receptor communication analysis R package, by taking the “to” genes in the file at https://zenodo.org/record/3260758/files/lr_network.rds. Secondly, we collected receptors by the keyword “receptor” from the UniProt database^33^ at https://www.uniprot.org/uniprot/?query=name%3Areceptor+reviewed%3Ayes+organism%3A%22Homo+sapiens+%28Human%29+%5B9606%5D%22&sort=score. The receptors from these two sources were united and further filtered by focusing on cell membrane proteins based on the cellular location annotation in the UniProt database. For cell surface transporters as the second category of sensor proteins, we noticed that the Transporter Classification Database (TCDB)^28-32^ contains a comprehensive list of transporter proteins. Thus, transporter gene names were collected from TCDB at https://www.tcdb.org/hgnc_explore.php. For nuclear receptors as the third category of sensor proteins, the Nuclear Receptor Signaling Atlas (NURSA)^34^ in dkNET project at https://dknet.org/data/source/nif-0000-03208-6/search was used for collecting names of known nuclear receptor names. After having the names of metabolites and sensors, text-mining based on PubMed abstracts were performed. Firstly, 711,272 PubMed abstracts were obtained from https://eutils.ncbi.nlm.nih.gov/entrez/eutils/esearch.fcgi?db=pubmed&term=%28metabolism%5BTitle%2FAbstract%5D%29&retmax=15000000. We started from those abstracts as they were related to metabolism. Such strategy helped us to narrow down to a subset of abstracts, thus reduced the time of text-mining compared to going through all the abstracts in the PubMed. Secondly, the context of publication titles, MeSH words, and abstracts was downloaded for those PMID. Thirdly, potential combinations of metabolite and sensor names was checked in each of the collected context of publication titles, MeSH words, and abstracts. The PMID, metabolite names, and sensor names were recorded if a pair of metabolite name and sensor name were co-mentioned in the same sentence in the context of publication titles, MeSH words, or abstracts. Next, the collected pairs of metabolite-sensor partners together with the PMID and evidence in the context of publication title, MeSH words, or abstracts were subjected to a manual curation.

Besides the text-mining, we also manually collected metabolite-sensor partners from other well-known databases such as the HMDB^20-23^, Recon2^35^, GPCRdb^37, 59^, Wikipedia, and GeneCards^36^. For HMDB, we collected metabolite and transporter pairs if the metabolite associated protein were annotated as a transporter and the protein belongs to a type of cell membrane protein. For Recon2, since metabolites were further annotated by cellular localization such as [e] for extracellular and [c] for cytosol, we specifically collected the reaction associated genes if the reaction happened by transporting the same metabolite from extracellular [e] into cytosol [c]. However, cell surface receptors and nuclear receptors were not included in both HMDB^20-23^ and Recon2^35^. Therefore, pairing of metabolite and cell surface transporter were mainly focused on those from these two databases. We additionally collected metabolite and cell surface receptor partners from GPCRdb^37, 38^ which is a database for G protein-coupled receptors (GPCR) basic annotation and their ligands. The metabolite ligand of the GPCR protein were collected from the webpage at https://gpcrdb.org/ligand/statistics if the metabolite is an extracellular metabolite. Additionally, one pair of metabolite-sensor partner, pyruvic acid and SLC16A11, was collected from the Gene Card database by reading the description of *SLC1611* at https://www.genecards.org/cgi-bin/carddisp.pl?gene=SLC16A11. Next, we manually collected metabolite and nuclear receptor partners from a Wikipedia page at https://en.wikipedia.org/wiki/Nuclear_receptor#Ligands.

All the metabolite-sensor partners were then further manually curated by at least three curators separately. Notably, the species and cell type were not restricted during the process of computational text-mining and manual collection, although the metabolite enzymes and sensor names were obtained mostly from databases for human. To generate a collection of metabolite-sensor partners for mouse, the metabolite enzyme genes and metabolite sensor genes were mapped to mouse genes based on the homology between human and mouse.

### Inferring metabolite presence in a cell based on expression of enzymes

Estimation of metabolite presence is a key step in MEBOCOST algorithm. Although several software, such as scFBA, scFEA, and COMPASS have been reported to predict metabolic flux rate by scRNA-seq data, none of them were specifically designed for estimating the metabolite distribution (Supplementary Figure 1C). Although the flux balance result in scFEA and the uptake reaction result in COMPASS can be treated as metabolite abundance as a by-product of the flux rate prediction, those tools still have limitations to be fully integrated into MEBOCOST. For example, scFEA predicts less than 100 metabolites, and COMPASS algorithm was computation-intensive so that it takes more than 1,000 seconds to run the prediction for a metabolite in 1,000 cells (Supplementary Figure 1E). Therefore, we tried to identify a method that can predict the metabolite abundance in a cell by scRNA-seq data with much less running time but without losing the accuracy when compared to the existing tools.

Although we did not incorporate the running of scFEA and COMPASS into the running of MEBOCOST, the development of such tools by other groups showed the power of metabolome modeling based on the RNA expression level of metabolic enzymes. We reasoned that the level of a given metabolite in a cell should be dynamically influenced by two types of metabolic reactions. Some reactions generate the metabolites as products; thus, the expression of the enzymes should correlate with the accumulation of the metabolites. Some other reactions may take the metabolites as substrates. The happening of such reactions will convert the given metabolites into other ones, so the enzyme expression should correlate with the depletion of the given metabolites. Therefore, we might be able to estimate the likelihood of presence of a given metabolite in a cell by the average expression of the enzymes that generate the metabolite as a product after subtracting the average expression of the enzymes that take the metabolite as a substrate. Our estimation of metabolite presence based on enzyme gene expression was formulated as:

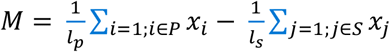

where *M* is the estimated presence of a given metabolite in a cell. *P* is a set of enzyme genes that are associated with the accumulation of the metabolite. *i* is the *i*^th^ enzyme gene in the *P. l*_*p*_ is the total number of enzyme genes in the *P*. In addition, *S* is a set of enzyme genes that are associated with the depletion of the metabolite. *j* is the *j*^th^ enzyme gene in the *S. l*_*s*_ is the total number of enzyme genes in the *S*. *x*_*i*_ and *x*_*j*_ are the expression level of enzyme *i* and *j* in a cell. MEBOCOST applied this formula to calculate the relative presence of each metabolite in each cell by scRNA-seq data.

To test the proposed method together with scFEA and COMPASS, we downloaded matched metabolomics and RNA-seq data of 928 cancer cell lines from the CCLE project^25, 26^. The metabolomics data was downloaded by https://depmap.org/portal/download/api/download?file_name=ccle%2Fccle_2019%2FCCLE_metabolomics_20190502.csv&bucket=depmap-external-downloads. The RNA-seq data was downloaded from DepMap data portal at https://depmap.org/portal/download/ (by the file name CCLE_expression.csv). Next, we applied scFEA, COMPASS, and our method on the expression matrix of the RNA-seq data to estimate the metabolite level in each cell line. In addition to taking the average (arithmetic mean) of the enzyme expression, we also calculated the geometric mean of the enzyme gene expression in the estimation for a comparison. To evaluate the similarity between the estimation and the real detection of metabolomics, the Jaccard index between the results of estimation and detection was calculated for each metabolite. To calculate the Jaccard index, cell lines that showed the presence of each metabolite were obtained for each estimation method as well as the real metabolomics data. Among the 910 cell lines, a metabolite was recognized as present in a cell line if the estimated or detected value of a given metabolite in that cell line is above the mean value of all the cell lines. The subset of cell lines that showed the presence of a metabolite based on the estimation data can be noted as A, and the subset of cell lines that showed the presence of a metabolite based on the metabolomics data was noted as B. The Jaccard index score *J* was calculated by

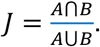

Interestingly, the Jaccard index value of the methods that are based on mean expression of enzyme genes showed quiet comparable result with scFEA and slightly better result than COMPASS (Supplementary Figure 1D). However, the methods that are based on arithmetic mean of enzyme gene expression appeared to be substantially faster than scFEA and COMPASS (Supplementary Figure 1E).

### Calculation of metabolic communication scores

Given the inferred metabolites for individual cell types and the prior knowledge of metabolite-sensor partners, MEBOCOST computed the metabolite-sensor communication score for each metabolite-sensor partner between each pair of cell types. For a pair of cell types *i* and *j*, and for a metabolite *m* and its sensor 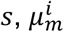 denotes the mean metabolite abundance in cell type *i*, and 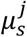 denotes the mean expression level of sensor gene in cell type *j*. The communication score *S*_*c*_ was computed as:

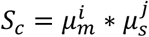

To evaluate the statistical significance of *S*_*c*_, we performed permutation testing^60^ by shuffling cell labels of the scRNA-seq data. For each metabolite-sensor partner between each pair of cell types, the same method was applied to calculate the communication score for each set of shuffled scRNA-seq data. We repeated this procedure 1,000 times to compute 1,000 communication scores as a statistical null distribution for each metabolite-sensor partner between each pair of cell types. Based on the *null* distribution, a *p* value was computed. The *p* values of all communication scores for a dataset were subjected to a false discovery rate (FDR) correction by the Benjamini-Hochberg method^27^. Furthermore, MEBOCOST provide three parameters, including “cutoff_exp”, “cutoff_met”, and “cutoff_prop”, for users to define expressed metabolite and sensor in cell populations. “cutoff_exp” and “cutoff_met” were cutoffs to define expressed metabolite and sensor in each cell, respectively. “cutoff_prop” was a cutoff to define the proportion of cells that expressed a metabolite and a sensor in a cell group (e.g. cell type). By default, “cutoff_exp” and “cutoff_met” were determined by taking 25% percentile value of all sensor expression and all metabolite presence in all cells, respectively. The default “cutoff_prop” was set to 0.15 which means that at least a 15% of total cells in the cell group expressed the metabolite and sensor. The p value and FDR of any communications with values lower than the three cutoffs will be converted to 1.

### Single cell RNA-seq data processing

The single cell RNA-seq data of mouse brown adipose tissue from TN, RT, Cold2, and Cold7 was generated by 10X Genomics platform. Raw data was deposited at NCBI GEO database under accession number GSE160585^39^. Cellranger was applied to map the raw sequence to mouse reference genome (mm10) and obtain read count over each gene, with r1-length parameter was set to 26 and other parameters were the default values. Next, the data processing, including data normalization, dimension reduction, clustering, and visualization of gene expression were performed using Scanpy^61^. Cells were filtered to have at least 800 UMIs and 400 detected genes. Genes were filtered to be at least detected in 10 cells. To reduce the doublet effect, cells were removed if the total UMIs were more than 50,000 or the number of detected genes is greater than 7,500. The number of nearest neighbors was set to 10 in the Scanpy “find neighbor” function. The top 40 principal components were included in clustering and UMAP analysis^62^. The visualization of the clusters was performed by the UMAP method. Cell annotation was done based on the cell type marker genes collected from PanglaoDB^63^ at https://panglaodb.se/markers.html.

### Evaluation of MEBOCOST running time and memory usage

We tested MEBOCOST on the BAT scRNA-seq data from mouse housed at cold temperature for 2 days (Cold2)^39^. The dataset contains 33,470 cells and 20 cell types. To run MEBOCOST, the parameters “cutoff_exp” and “cutoff_met” were set to 0.866 and 0.141, respectively. The two values were determined by taking the 25% percentile value of all sensor expression and all metabolite presence in all cells in our BAT scRNA-seq dataset, respectively. The parameter “cutoff_prop” was set to 0.15 to require at least 15% of cells expressed the metabolites and sensors in the cell type. All the jobs were run on same computing server with 8 cores. The running time were recorded by the python “time” module, and the peak memory usage were also recorded by the python “tracemalloc” module.

### Evaluation of MEBOCOST performance stability

We reasoned that a good algorithm should be less likely to be influenced by the sequencing depth and the total cell number in scRNA-seq data. Therefore, we compared the prediction results of subsampling datasets with the result of original dataset in two aspects. First, we compared the total number of significant communication events between subsampling datasets and original datasets. The communications were deemed significant at a False Discovery Rate (FDR) of 0.05. Second, the similarity between the prediction result of subsampling datasets and the original dataset was evaluated. Two overlapping percentages were calculated to measure such similarity. One is the number of overlapped communications between subsampled datasets and the original dataset divided by the number of communications in the subsampled data (recaptured over total detected). The other one is the number of overlapped communications between subsampled datasets and the original dataset divided by the number of communications in the original dataset (recaptured over original).

To test the effect of sequencing noise on the performance of MEBOCOST, we added in silico noise to scRNA-seq dataset with a series of noise rates. Then noise rate was calculated as number of random reads divided by the number or scRNA-seq reads in the original dataset. Each noised dataset was generated by introducing random reads into the Cold2 scRNA-seq data. The detailed procedures mainly include three steps. First, the total number of noise reads was calculated by multiplying the total read count of Cold2 scRNA-seq data by the given noise rate. Second, data points were randomly selected from the cell-by-gene count matrix. Third, the original read count in the selected data point was added by one. Four, the second and third steps were iterated and stopped until all noise were added to reach the noise rate. The newly generated cell-by-gene count matrix of noised datasets was then subjected to normalization and log transformation by the Scanpy package. Next, MEBOCOST was applied on each noised dataset to detect metabolite-sensor communications.

### Identification of the most cold-sensitive communication events

All the cells of BAT scRNA-seq data from TN, RT, Cold2, and Cold7 were pooled and grouped by cell types and conditions when running MEBOCOST. Communications were deemed as significant if FDR is less than 0.05. Index of dispersion (IOD) was calculated for communication score across the four conditions. Given a sender-metabolite-sensor-receiver combination, the IOD was defined by:

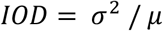

where *σ*^2^ is the variance of the four scores of the communication under the four conditions, *µ* is the mean of the four scores. Sender-metabolite-sensor-receiver communications included in this cold-sensitive communication analysis were defined as significant at least in one of the four conditions. The IOD scores of communications were ranked from high to low, and the 100 top-ranked were defined as cold-sensitive communications.

## Code availability and data availability

All the algorithm in MEBOCOST were implemented by Python. The collected enzyme and sensor genes of extracellular metabolites, source code of MEBOCOST, and the detailed instruction of usage were available at https://github.com/zhengrongbin/MEBOCOST. The single cell RNA-seq data of mouse brown adipose tissue is available at NCBI GEO under the accession number GSE160585.

## Acknowledgements

This project is supported in part by R01GM125632 (Kaifu Chen), 1R01HL148338 (Kaifu Chen), R01DK132469 (Yu-Hua Tseng), and R01HL155632 (Lili Zhang).

## Author Contributions

K. C. conceived the project. R. Z. developed the algorithm, curated the metabolite-sensor partners, and performed the computational analysis. K. C., Y. Z., and T. T. curated the metabolite-sensor partners. K.C., Y.H.T., R.Z., and L.Z analyzed and interpreted the results. R. Z. and K. C. drafted and edited the manuscript. All authors read and edited the manuscript.

## Abbreviations

EC: Endothelial cells
CD8T: CD8^+^ T lymphocytes
Activated CD8T: Activated CD8^+^ T lymphocytes
CD4T: CD4^+^ T lymphocytes
Erythroid-like: Erythroid-like and erythroid precursor cells
NK: Natural killer cells
MSC: Myelinating Schwann cells
NMSC: Non-myelinating Schwann cells
Treg: Regulatory T cells
VSM: Vascular smooth muscle cells
ILC2s: Type 2 innate lymphoid cells
BAT: Brown adipose tissue
TN: Thermoneutrality
RT: Room temperature
Cold2: Cold temperature for 2 days
Cold7: Cold temperature for 7 days

---

**Supplementary Figure 1.**
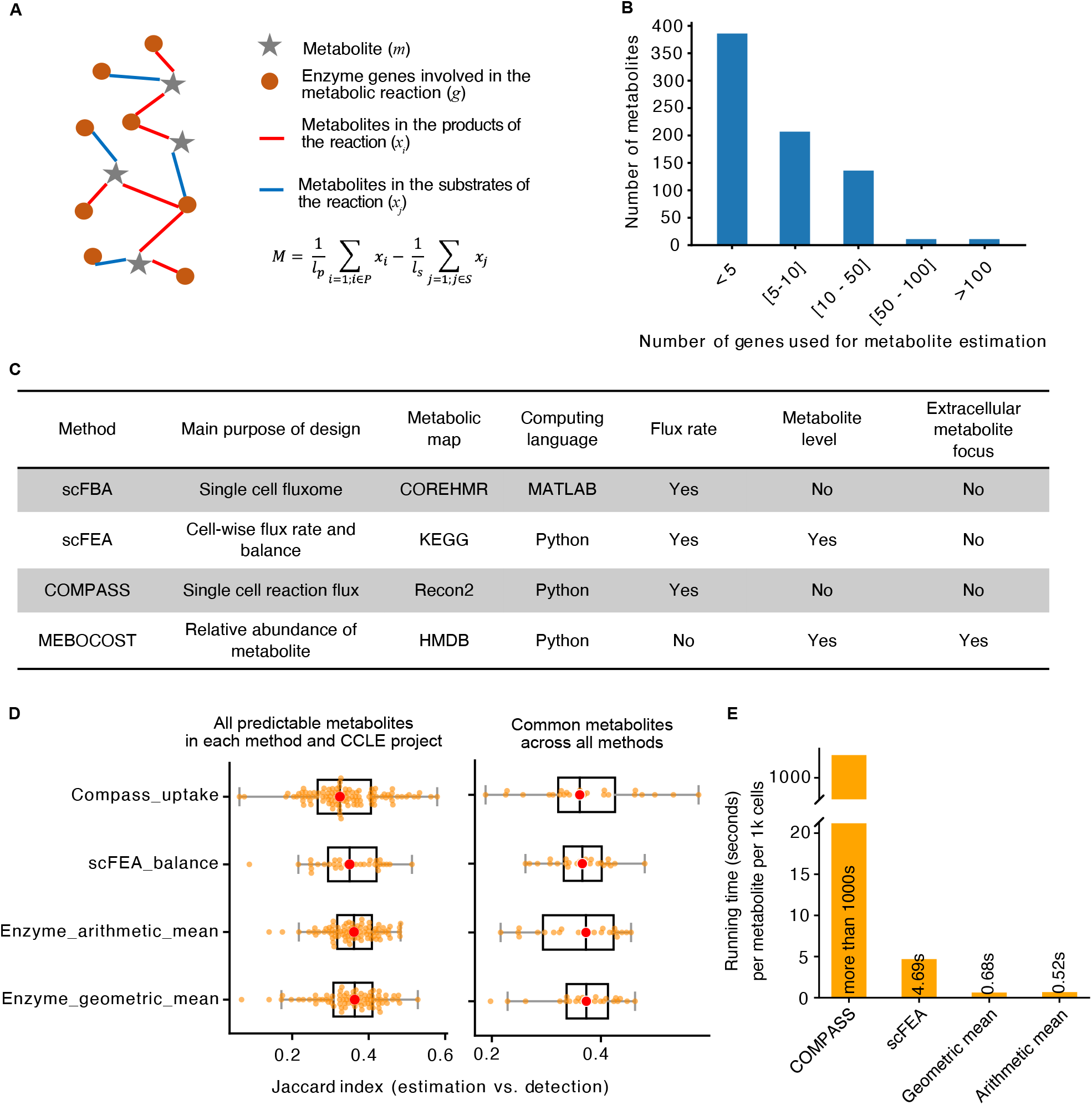
Estimation of metabolite presence based on RNA expression of metabolic enzymes. **A**. The schematic plot showed for metabolite presence estimation. **B**. Number of metabolites plotted against number of metabolic enzymes per metabolite. **C**. A summary of tools for estimating metabolite presence. **D**. Boxplot showing accuracy for inference of metabolite presence. RNA-seq data from the CCLE dataset were used for estimation; the matched metabolomics profiles of same samples in the CCLE dataset were used as the ground truth for detection of metabolites. Because some metabolites are not covered by all these methods, the metabolites covered by individual method were used for analysis in the left panel, where the subset of metabolites covered by all methods were used for analysis in the right panel. **E**. Bar plot showing the running time of individual methods for inference of metabolite presence.

**Supplementary Figure 2.**
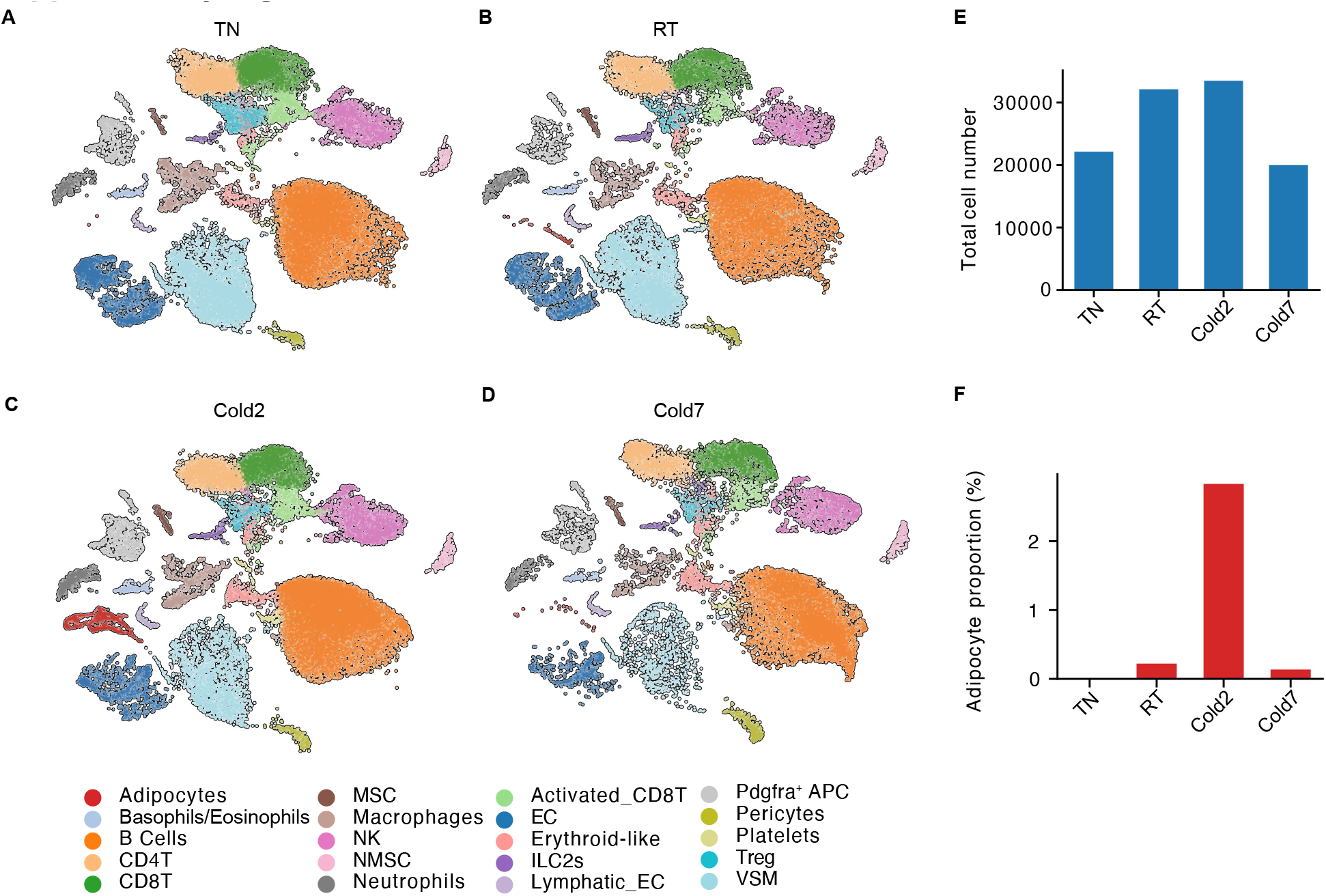
Single cell RNA-seq data of mouse adipose tissue. **A-D**. UMAP visualization of the scRNA-seq data from four conditions, including TN (30 °C for a week) (A), RT (room temperature) (B), Cold2(4 °C for 2 days) (C), and Cold7 (4 °C for 7 days) (D). **E**. The total number of cells in the scRNA-seq data from the four conditions. **F**. The proportion of differentiating adipocytes across the four conditions.

**Supplementary Figure 3.**
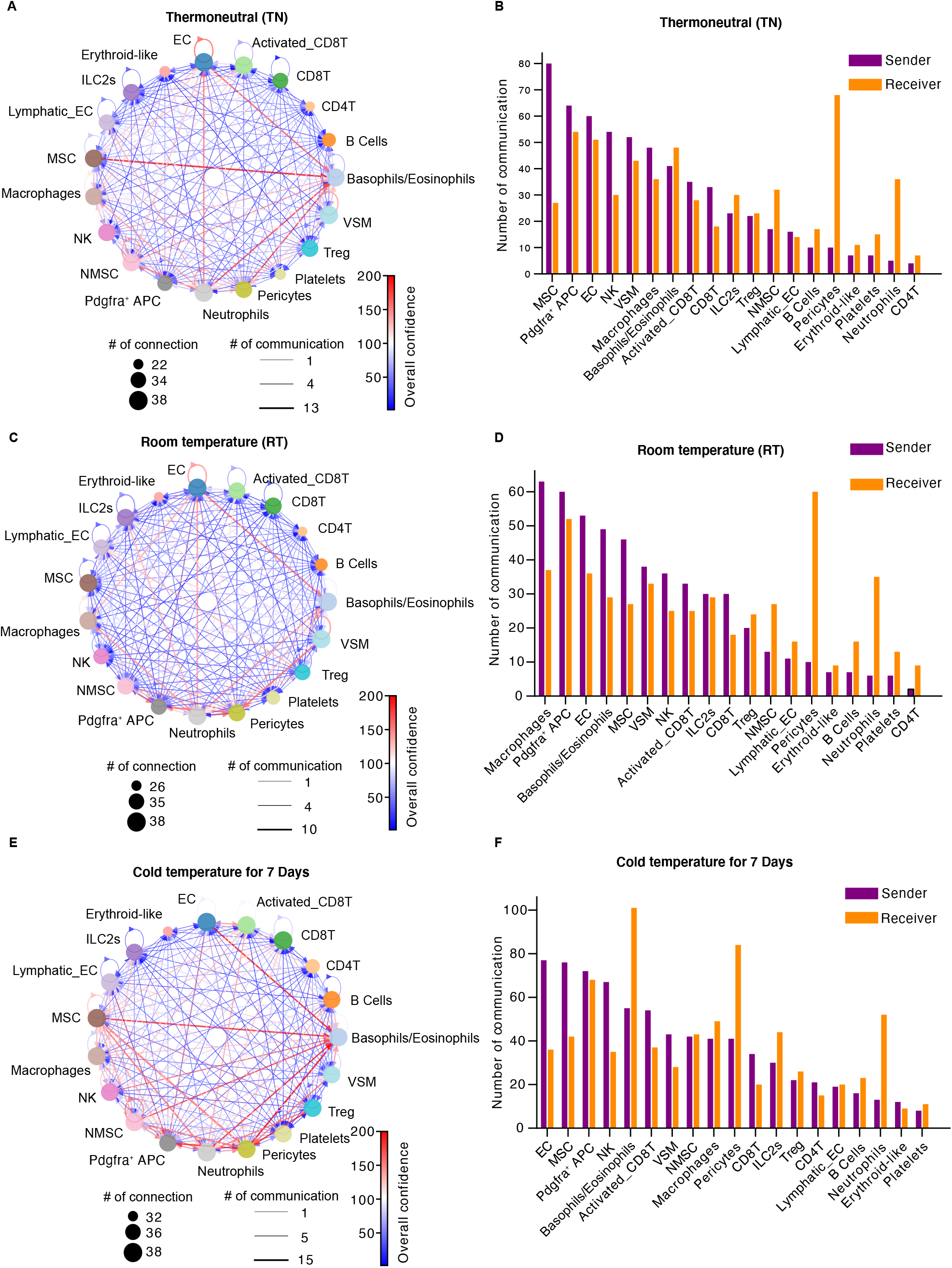
The metabolite-sensor communications detected by MEBOCOST in BAT of mouse from the TN, RT, and Cold7 conditions. **A, C, E**. Network plot showing cell-to-cell metabolite-sensor communications detected by MEBOCOST for mouse BAT at the TN (A), RT (C), and Cold7 (E) conditions. Each dot was a cell type. The size of dot for each cell type represented the number of communications with the other cell types. Each arrow line represented the communication from a sender cell type to a receiver cell type. The line width indicated the number of metabolite-sensor communications between the sender and receiver cell types. Line color showed the overall communication score which was calculated by the sum of −log10(FDR) of all metabolite-sensor communications between the sender and receiver types. **B, D, F**. Bar plot to show the number of detected communications with each cell type as sender cells or receiver cells at the TN (B), RT (D), and Cold7 (F) conditions.

## References

1. Song, D., Yang, D., Powell, C.A. & Wang, X. Cell-cell communication: old mystery and new opportunity. Cell Biol Toxicol 35, 89–93 (2019).

2. Lafontan, M. Fat cells: afferent and efferent messages define new approaches to treat obesity. Annu Rev Pharmacol Toxicol 45, 119–146 (2005).

3. Roy, S., Kim, D. & Lim, R. Cell-cell communication in diabetic retinopathy. Vision Res 139, 115–122 (2017).

4. Tirziu, D., Giordano, F.J. & Simons, M. Cell communications in the heart. Circulation 122, 928–937 (2010).

5. AlMusawi, S., Ahmed, M. & Nateri, A.S. Understanding cell-cell communication and signaling in the colorectal cancer microenvironment. Clin Transl Med 11, e308 (2021).

6. Armingol, E., Officer, A., Harismendy, O. & Lewis, N.E. Deciphering cell-cell interactions and communication from gene expression. Nat Rev Genet 22, 71–88 (2021).

7. Jin, S. et al. Inference and analysis of cell-cell communication using CellChat. Nat Commun 12, 1088 (2021).

8. Efremova, M., Vento-Tormo, M., Teichmann, S.A. & Vento-Tormo, R. CellPhoneDB: inferring cell-cell communication from combined expression of multi-subunit ligand-receptor complexes. Nat Protoc 15, 1484–1506 (2020).

9. Martins Conde Pdo, R., Sauter, T. & Pfau, T. Constraint Based Modeling Going Multicellular. Front Mol Biosci 3, 3 (2016).

10. Dias, A.S., Almeida, C.R., Helguero, L.A. & Duarte, I.F. Metabolic crosstalk in the breast cancer microenvironment. Eur J Cancer 121, 154–171 (2019).

11. Wagner, A. et al. Metabolic modeling of single Th17 cells reveals regulators of autoimmunity. Cell 184, 4168–4185 e4121 (2021).

12. Alghamdi, N. et al. A graph neural network model to estimate cell-wise metabolic flux using single-cell RNA-seq data. Genome Res 31, 1867–1884 (2021).

13. Damiani, C. et al. Integration of single-cell RNA-seq data into population models to characterize cancer metabolism. PLoS Comput Biol 15, e1006733 (2019).

14. Wang, Y.P. & Lei, Q.Y. Metabolite sensing and signaling in cell metabolism. Signal Transduct Target Ther 3, 30 (2018).

15. Liu, S., Alexander, R.K. & Lee, C.H. Lipid metabolites as metabolic messengers in inter-organ communication. Trends Endocrinol Metab 25, 356–363 (2014).

16. Monelli, E. et al. Angiocrine polyamine production regulates adiposity. Nat Metab (2022).

17. Schloss, M.J. et al. B lymphocyte-derived acetylcholine limits steady-state and emergency hematopoiesis. Nature Immunology (2022).

18. Li, H. et al. The allergy mediator histamine confers resistanceto immunotherapy in cancer patients via activationof the macrophage histamine receptor H1. Cancer Cell (2021).

19. Sun, H. & Wang, Y. A new branch connecting thermogenesis and diabetes. Nat Metab 1, 845–846 (2019).

20. Wishart, D.S. et al. HMDB 4.0: the human metabolome database for 2018. Nucleic Acids Res 46, D608–D617 (2018).

21. Wishart, D.S. et al. HMDB 5.0: the Human Metabolome Database for 2022. Nucleic Acids Res 50, D622–D631 (2022).

22. Wishart, D.S. et al. HMDB 3.0--The Human Metabolome Database in 2013. Nucleic Acids Res 41, D801–807 (2013).

23. Wishart, D.S. et al. HMDB: the Human Metabolome Database. Nucleic Acids Res 35, D521–526 (2007).

24. Tonn, M.K., Thomas, P., Barahona, M. & Oyarzun, D.A. Computation of Single-Cell Metabolite Distributions Using Mixture Models. Front Cell Dev Biol 8, 614832 (2020).

25. Barretina, J. et al. The Cancer Cell Line Encyclopedia enables predictive modelling of anticancer drug sensitivity. Nature 483, 603–607 (2012).

26. Li, H. et al. The landscape of cancer cell line metabolism. Nat Med 25, 850–860 (2019).

27. Hochberg, Y.B.a.Y. Controlling the False Discovery Rate: a Practical and Powerful Approach to Multiple Testing. J. R. Statist. Soc 57, 289–300 (1995).

28. Saier, M.H. et al. The Transporter Classification Database (TCDB): 2021 update. Nucleic Acids Res 49, D461–D467 (2021).

29. Saier, M.H., Jr., Reddy, V.S., Tamang, D.G. & Vastermark, A. The transporter classification database. Nucleic Acids Res 42, D251–258 (2014).

30. Saier, M.H., Jr. et al. The Transporter Classification Database (TCDB): recent advances. Nucleic Acids Res 44, D372–379 (2016).

31. Saier, M.H., Jr., Tran, C.V. & Barabote, R.D. TCDB: the Transporter Classification Database for membrane transport protein analyses and information. Nucleic Acids Res 34, D181–186 (2006).

32. Saier, M.H., Jr., Yen, M.R., Noto, K., Tamang, D.G. & Elkan, C. The Transporter Classification Database: recent advances. Nucleic Acids Res 37, D274–278 (2009).

33. UniProt, C. UniProt: the universal protein knowledgebase in 2021. Nucleic Acids Res 49, D480–D489 (2021).

34. Becnel, L.B. et al. Nuclear Receptor Signaling Atlas: Opening Access to the Biology of Nuclear Receptor Signaling Pathways. PLoS One 10, e0135615 (2015).

35. Thiele, I. et al. A community-driven global reconstruction of human metabolism. Nat Biotechnol 31, 419–425 (2013).

36. Safran, M. et al. GeneCards Version 3: the human gene integrator. Database (Oxford) 2010, baq020 (2010).

37. Kooistra, A.J. et al. GPCRdb in 2021: integrating GPCR sequence, structure and function. Nucleic Acids Res 49, D335–D343 (2021).

38. Pandy-Szekeres, G. et al. The G protein database, GproteinDb. Nucleic Acids Res 50, D518–D525 (2022).

39. Shamsi, F. et al. Vascular smooth muscle-derived Trpv1(+) progenitors are a source of cold-induced thermogenic adipocytes. Nat Metab 3, 485–495 (2021).

40. Lee, P. et al. Temperature-acclimated brown adipose tissue modulates insulin sensitivity in humans. Diabetes 63, 3686–3698 (2014).

41. Peres Valgas da Silva, C., Hernandez-Saavedra, D., White, J.D. & Stanford, K.I. Cold and Exercise: Therapeutic Tools to Activate Brown Adipose Tissue and Combat Obesity. Biology (Basel) 8 (2019).

42. van Marken Lichtenbelt, W.D. et al. Cold-activated brown adipose tissue in healthy men. N Engl J Med 360, 1500–1508 (2009).

43. McMillan, A.C. & White, M.D. Induction of thermogenesis in brown and beige adipose tissues: molecular markers, mild cold exposure and novel therapies. Curr Opin Endocrinol Diabetes Obes 22, 347–352 (2015).

44. Whitehead, A. et al. Brown and beige adipose tissue regulate systemic metabolism through a metabolite interorgan signaling axis. Nat Commun 12, 1905 (2021).

45. Singh, R., Braga, M. & Pervin, S. Regulation of brown adipocyte metabolism by myostatin/follistatin signaling. Front Cell Dev Biol 2, 60 (2014).

46. Leiria, L.O. et al. 12-Lipoxygenase Regulates Cold Adaptation and Glucose Metabolism by Producing the Omega-3 Lipid 12-HEPE from Brown Fat. Cell Metab 30, 768–783 e767 (2019).

47. Kim, J. et al. Eicosapentaenoic Acid Potentiates Brown Thermogenesis through FFAR4-dependent Up-regulation of miR-30b and miR-378. J Biol Chem 291, 20551–20562 (2016).

48. Herz, C.T. & Kiefer, F.W. The Transcriptional Role of Vitamin A and the Retinoid Axis in Brown Fat Function. Front Endocrinol (Lausanne) 11, 608 (2020).

49. Fenzl, A. et al. Intact vitamin A transport is critical for cold-mediated adipose tissue browning and thermogenesis. Mol Metab 42, 101088 (2020).

50. Yoo, H., Antoniewicz, M.R., Stephanopoulos, G. & Kelleher, J.K. Quantifying reductive carboxylation flux of glutamine to lipid in a brown adipocyte cell line. J Biol Chem 283, 20621–20627 (2008).

51. Okamatsu-Ogura, Y. et al. UCP1-dependent and UCP1-independent metabolic changes induced by acute cold exposure in brown adipose tissue of mice. Metabolism 113, 154396 (2020).

52. Shamsi, F., Wang, C.H. & Tseng, Y.H. The evolving view of thermogenic adipocytes - ontogeny, niche and function. Nat Rev Endocrinol 17, 726–744 (2021).

53. Hammel, J.H. & Bellas, E. Endothelial cell crosstalk improves browning but hinders white adipocyte maturation in 3D engineered adipose tissue. Integr Biol (Camb) 12, 81–89 (2020).

54. Garcia-Alonso, L. et al. Single-cell roadmap of human gonadal development. Nature (2022).

55. Jakobsson, J.E.T., Spjuth, O. & Lagerstrom, M.C. scConnect: a method for exploratory analysis of cell-cell communication based on single cell RNA sequencing data. Bioinformatics (2021).

56. Robinson, J.L. et al. An atlas of human metabolism. Sci Signal 13 (2020).

57. Koppula, P., Zhuang, L. & Gan, B. Cystine transporter SLC7A11/xCT in cancer: ferroptosis, nutrient dependency, and cancer therapy. Protein Cell 12, 599–620 (2021).

58. Browaeys, R., Saelens, W. & Saeys, Y. NicheNet: modeling intercellular communication by linking ligands to target genes. Nat Methods 17, 159–162 (2020).

59. Husted, A.S., Trauelsen, M., Rudenko, O., Hjorth, S.A. & Schwartz, T.W. GPCR-Mediated Signaling of Metabolites. Cell Metab 25, 777–796 (2017).

60. Moore, J.H. Bootstrapping, permutation testing and the method of surrogate data. Phys Med Biol 44, L11–12 (1999).

61. Wolf, F.A., Angerer, P. & Theis, F.J. SCANPY: large-scale single-cell gene expression data analysis. Genome Biol 19, 15 (2018).

62. Becht, E. et al. Dimensionality reduction for visualizing single-cell data using UMAP. Nat Biotechnol (2018).

63. Franzen, O., Gan, L.M. & Bjorkegren, J.L.M. PanglaoDB: a web server for exploration of mouse and human single-cell RNA sequencing data. Database (Oxford) 2019 (2019).

